# Diet-Associated Differences in the Rumen Microbiome and Resistome of Bangladeshi Cattle

**DOI:** 10.64898/2026.04.27.721148

**Authors:** Sunjukta Ahsan, Mohammad Nurul Islam, Nur A. Hasan, Michael Netherland, Moumita Chakrabarti, Faiza Noor, Ellin Ferdous Mohona

**Affiliations:** Department of Microbiology, University of Dhaka, Bangladesh; Department of Botany, University of Dhaka, Bangladesh; Arius Bioscience, Inc., Maryland, USA; Department of Microbiology, BRAC University

## Abstract

Diet influences the composition, diversity, and functional capacity of the cattle gut microbiome. However, the extent to which feeding practices affect the microbial community and resistome under real-world conditions remains poorly understood, particularly in low- and middle-income settings. Here, we applied metagenomics to fecal samples from Bangladeshi cattle fed either a natural or a mixed diet to examine differences in microbial composition, functional potential, and resistome associated with feed type. Natural-fed cattle harbored higher microbial diversity and distinct bacterial phyla, including Bacteroidota, Campylobacteriota, and Mycoplasmatota. *Acinetobacter, Aliarcobacter, Comamonas, Dysosmobacter*, and *Sharpea* were enriched in natural-fed cattle, whereas *Anaerotignum, Aristaeella, Oscillibacter*, and *Clostridium* were more abundant in the mixed-fed group. Notably, the emerging zoonotic genus *Aliarcobacter* was detected in the natural-fed cohort. Alpha diversity analysis showed higher richness and evenness in natural-fed cattle, and a clear separation between dietary groups in beta diversity analysis (PERMANOVA, p = 0.01). Differential analysis identified *Oscillibacter ruminantium* as a biomarker of natural feeding, while *Succinivibrio faecicola* and *Anaerovibrio slackiae* for mixed feeding. Resistome profiles demonstrated clear differences. Mixed-fed cattle showed a consistent enrichment of tetracycline resistance genes, whereas the natural-fed group displayed a more variable resistome. Functional analysis suggested diet-associated differences in metabolic potential, with glutathione metabolism enriched in natural-fed cattle (*p*<0.05) and bile secretion and fatty acid metabolism moderately enriched in the mixed-fed group. These findings indicate that feeding practices are associated with differences in rumen microbial communities and resistome profiles in Bangladeshi cattle, providing baseline insights into microbiome–resistome relationships under field conditions.

## Introduction

Ruminants are a major source of meat and milk derived from plant biomass (Eisler et al., 2014). The rumen microbiome plays a central role in converting this biomass into usable energy and improving overall feed efficiency (O’Hara et al., 2020). It is responsible for breaking down low-quality feed into absorbable nutrients, providing approximately 70% of the energy required by the host animal (Mao et al., 2015). Microorganisms from multiple kingdoms, including Bacteria, Archaea, Protozoa, and Fungi, contribute to this process through proteolytic, fibrolytic, and lipolytic activities. The end products include volatile fatty acids, biohydrogenated lipids, and other metabolites, while Microbial Crude Protein (MCP) serves as an important source of protein for the host (Stewart et al., 2019; Alberdi et al., 2021).

The diversity and composition of the rumen microbiome are influenced by several factors, including feed efficiency (Eastridge, 2006), disease status (Zhao et al., 2025), and methane emission (Petra et al., 2014). In turn, microbial diversity affects host metabolism and overall physiological performance (Martínez-Álvaro et al., 2020; Artzi et al., 2017; Seshadri et al., 2018). External inputs such as antibiotics, feed additives, and probiotics can also alter the rumen microbial ecosystem, sometimes disrupting its stability (Chae et al., 2020; Dimroth and Schink, 1998; Leahy et al., 2022; Cholewinska et al., 2020).

Bacteria make up roughly half of the rumen microbial community, followed by protozoa, fungi, and methanogenic Archaea (Ransom et al., 2012). The dominant bacterial phyla are Firmicutes and Bacteroidetes, with Firmicutes generally associated with forage-based diets and Bacteroidetes more abundant in concentrate-fed animals (Driks, 2003; Liggenstoffer et al., 2010). These are followed by other groups such as Proteobacteria, Tenericutes, and Actinobacteria (Tropini et al., 2017). Within these communities, Prevotella species are particularly widespread due to their broad functional capabilities (Borrel et al., 2016; Donaldson et al., 2016). Despite variation across species, diets, and geographic regions, a core microbiome is typically maintained in the rumen (Kamke et al., 2016).

In Bangladesh, livestock feeding practices are largely based on locally available resources, including crop residues, grasses, and tree leaves, with limited use of concentrated feed (Huque and Sarker, 2014). At the same time, antimicrobials are often incorporated into feed to prevent disease and promote growth (Kirbis, 2007; Islam et al., 2016). While antibiotic use can influence the rumen microbiome, these effects are often transient and may vary depending on the production system (Lin et al., 2019). Under such field conditions, the combined effects of feeding practices and environmental exposure on microbial communities and antimicrobial resistance remain poorly understood.

In this study, we investigated how different feeding practices are associated with changes in the rumen microbiome and antimicrobial resistance profiles in Bangladeshi cattle. We applied shotgun metagenomic sequencing to fecal samples from cows fed either a natural grass-based diet or a mixed diet of grass and commercial feed. We aimed to characterize taxonomic composition, identify feed-associated microbial signatures, and examine patterns of antimicrobial resistance and functional potential. As part of this analysis, we also report the detection of the emerging zoonotic genus *Aliarcobacter* in Bangladeshi cattle. This pilot study provides baseline data to better understand microbiome–resistome relationships under real-world production conditions and may help inform feeding strategies and antimicrobial stewardship in livestock systems.

## Methods

### Sample collection

Fecal samples were collected from a total of 16 cows across various locations in Bangladesh. Based on farmer-reported feedback on feed types, two cohorts of eight animals each were sampled to represent (a) Natural-fed group from farms in the Rangpur and Jashore districts, and (b) Mixed-fed group from farms in the Sadarghat, Chankharpul, and Gazipur districts. In each case, fresh feces were aseptically collected from the cows and transferred into sterile falcon tubes. The samples were transported to the laboratory on ice and stored in a freezer at -20°C until further processing.

### DNA extraction and sequencing

Total genomic DNA was extracted from 200 mg of cow feces via a QIAmp Fast DNA Stool Kit (Qiagen, Germany). The purity of the DNA was monitored via a 1% agarose gel. Metagenomic shotgun sequencing of the DNA was performed by Arius Biosciences (Gaithersburg, MD, USA). The concentration of genomic DNA was measured via the Qubit Fluorometer dsDNA DNA quantification System (Thermo Fisher, USA). A total of 50 ng-1 µg of genomic DNA was used for library construction via the NEBNext® Ultra™ II FS DNA Library Prep Kit for Illumina®. Briefly, gDNA was enzymatically sheared, and the ends of the DNA fragments were repaired, 3’ adenylated, and ligated to adapters according to the manufacturer’s instructions. The resulting adapter-ligated libraries were PCR-amplified via the following protocol: an initial denaturation step at 98°C for 45 s; 5 cycles of denaturation (98°C, 15 s), annealing (60°C, 30 s) and extension (72°C, 30 sec); and a final elongation of 1 min at 72°C. The PCR products were removed from the reaction mixture with Mag-Bind RxnPure Plus magnetic beads (Omega Biotek, Norcross, GA). The libraries were quantified and qualified via D1000 ScreenTape on an Agilent 2200 TapeStation instrument. The libraries were normalized and pooled for multiplexed sequencing targeting 20 million reads per samples on an Illumina HiSeq X10 sequencer (Illumina, San Diego, CA, USA) via the paired-end 150 bp run format.

### Taxonomic and functional microbiome profiling

The profiling process started by surveying the potential presence of bacterial species for each raw metagenomic sample via Kraken2 (Wood, Lu, and Langmead, 2019) and a prebuilt core gene database (Chalita *et al*., 2020) containing k-mers (k=35) of reference genomes obtained from Arius DB. Fungi and Viral full genomes from NCBI’s refseq (https://www.ncbi.nlm.nih.gov/refseq/) were also added to the Kraken2 database. After a list of candidate species was acquired, a custom Bowtie2 (Langmead and Salzberg, 2012) database was built utilizing the core genes and genomes from the species identified during the first step. The raw sample was then mapped against the Bowtie2 database via the --very-sensitive option and a quality threshold of phred33. SAMtools (Li *et al*., 2009) was used to convert and sort the output bam files. The coverage of the mapped reads against the bam file was obtained via Bedtools (Quinlan and Hall, 2010). To avoid false positives, using an in-house script, we quantified all the reads that mapped to a given species only if the total coverage of their core genes (archaea, bacteria) or genome (fungi, virus) was at least 25%. Finally, species abundance was calculated from the total number of reads counted, and normalized species abundance was calculated from the total length of all the reference reads. For each sample, functional annotations were obtained by matching each read via DIAMOND (Buchfink, Xie, and Huson, 2014) against the KEGG database (Kanehisa *et al*., 2017). DIAMOND was executed via the blastx parameter, which converts each metagenomic read into multiple amino acid sequences by generating all six open reading frame variations and then matches it against the prebuilt KEGG database. If a read had multiple KEGG hits, the top hit was always used. After all the KEGG orthologs present were quantified, minpath (Ye and Doak, 2009) was used to predict the presence of KEGG functional pathways.

### Antimicrobial resistance gene profiling

Antibiotic resistance gene profiles were produced by using a prebuilt Bowtie2 (Langmead and Salzberg, 2012) database composed of NCBI’s National Database of Antibiotic-Resistant Organisms (NDARO, www.ncbi.nlm.nih.gov/pathogens/antimicrobial-resistance/) reference genes. Each read of the metagenome sample was mapped against these genes via Bowtie2 with the very sensitive option, and the output was then converted and sorted via SAMtools (Li *et al*., 2009). Finally, for each gene found, depth and coverage were calculated by using SAMtools’ mpileup script.

### Comparative metagenomic and statistical analyses

We used various comparative metagenomics and statistical methods to compare the microbiome composition, diversity, function, and resistome across different samples and feed types. For microbiome composition analysis, average relative abundances per phylum and genus per sample type were calculated and visualized via bar charts and sunburst charts. Alpha diversity was measured via Shannon (Shannon 1948) and Inverse Simpson (Simpson 1949). Beta diversity was analyzed via the pairwise Bray–Curtis distance index (Bray and Curtis, 1957), followed by principal coordinate analysis (PCoA). Pairwise PERMANOVAs, using 999 permutations, revealed significant differences between sample groups in the PCoA ordination. ALDEx2 result p-values were adjusted via the FDR method (“BH”) of the p.adjust() function in R (Benjamini and Hochberg, 1995). A matrix of pathway counts was produced from the functional pathways revealed by minpath by aggregating the individual KEGG ortholog read counts for each pathway. These raw counts were normalized via DESeq2’s median-of-ratios method (Love *et al*., 2014). All statistics for alpha diversity and pathway analyses were produced with the ‘ranksums’ function from SciPy.

## Results

### Sample Information

Samples were collected from five locations across Bangladesh. Fecal samples from naturally fed cows were collected from villages in Rangpur and Jassore districts. For mixed-fed cows, samples were collected from three farms located in Sadarghat, Chankharpul, and Gazipur in central Bangladesh (Dhaka division).

### Community Composition at Various Taxonomic Levels

At the phylum level, the microbiota of both groups were dominated by Actinomycetota, Bacillota, Pseudomonadota, and Thermodesulfobacteriota. Bacteriodota, Campylobacteriota, and Mycoplasmatota were detected only in the natural-fed samples. In contrast, *Candidatus* and *Spirochaetota* were observed only in the mixed-fed samples. Thermodesulfobacteriota were at higher relative abundance in natural-fed samples compared to mixed-fed samples (Figure 1).

**Figure 1.**
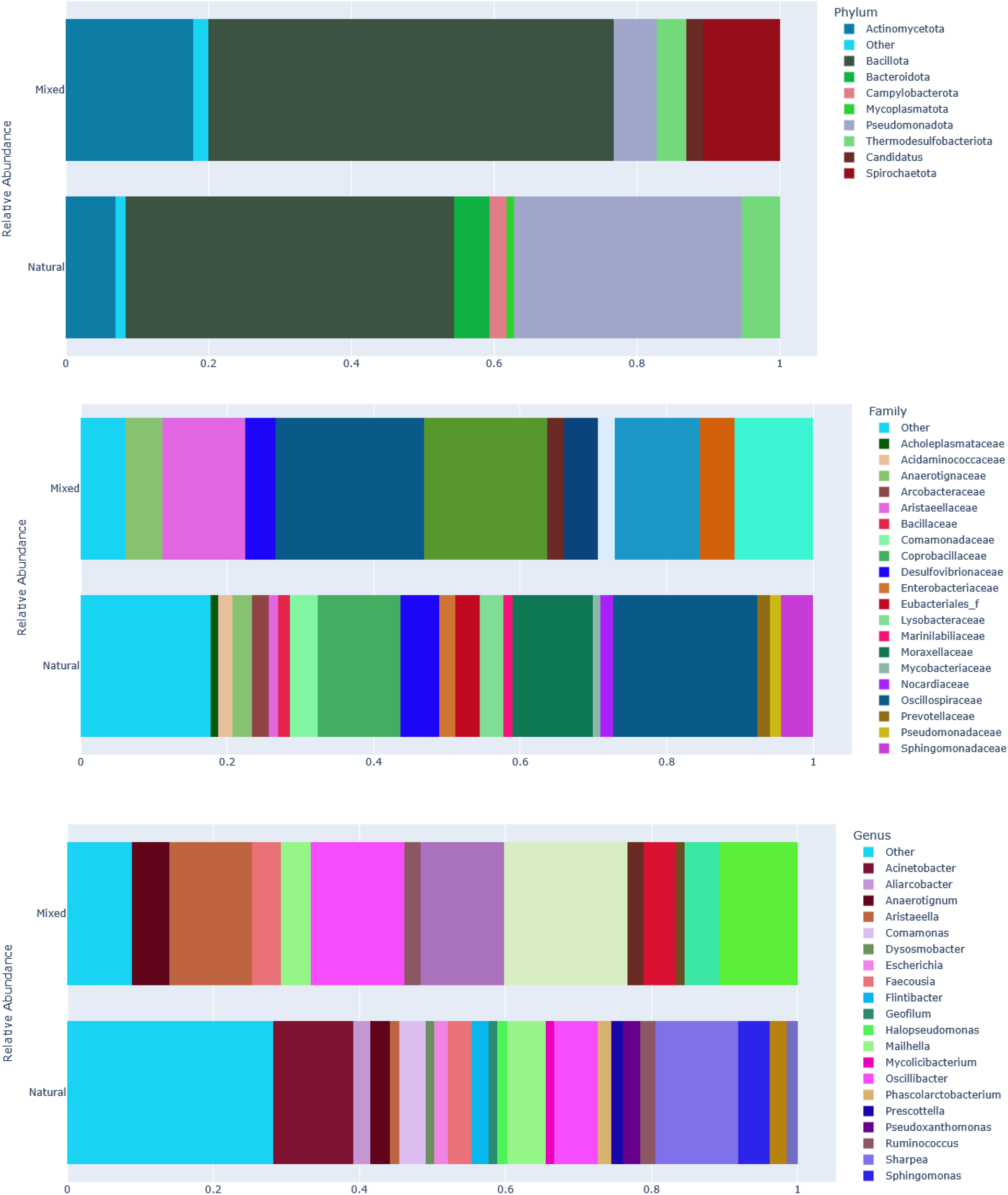
Relative abundance of rumen microbiota in natural- and mixed-fed cows at the phylum, family, and genus levels. Stacked bar plots illustrate dietary differences in microbial community composition across taxonomic ranks.

At the family level, greater variation was observed in the natural-fed samples than in the mixed-fed samples. Families including Coprobacteriaceae, Moraxellaceae, Acholeplasmataceae, Acidaminococcaceae, Arcobacteraceae, and Bacillaceae were detected only in the natural-fed group. The mixed-fed group showed comparatively lower diversity, with families such as Anaerotignaceae, Aristaeellaceae, and Oscilliospiraceae being detected in both groups (Figure 2).

**Figure 2.**
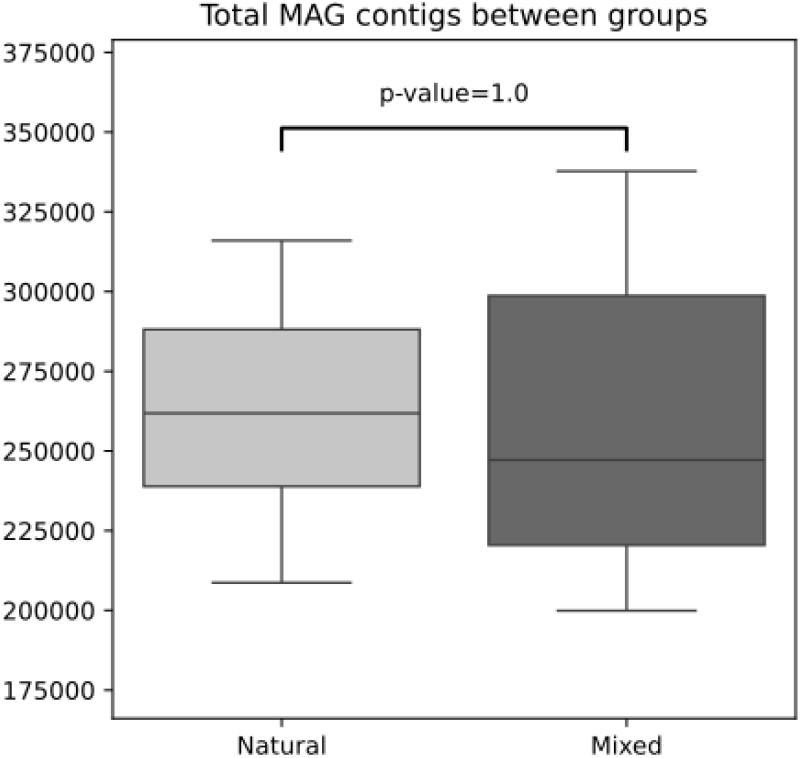
The distribution of metagenome-assembled genome (MAG) contig counts between dietary cohorts. The *p-value* from a statistical test comparing the two groups is indicated.

At the genus level, diversity remained consistently higher in the natural-fed group. Genera such as *Anaerotignum, Aristaeella*, and *Oscillibacter* were more abundant in the mixed-fed group. In contrast, *Acinetobacter, Aliarcobacter, Comamonas, Dysomobacter*, and *Sharpea* were enriched in the natural-fed group. Additional genera, including *Flintibacter, Geofilum, Halopseudomonas, Mycolicibacterium, Phascolarctobacterium*, and *Prescottella* were also detected in the natural-fed group at a moderate abundance.

### Metagenomic Assembled Genomes (MAGs)

Metagenomic assembly generated contig-level data; however, no high-quality metagenome-assembled genomes (MAGs) were recovered (Figure 2). Therefore, the results are presented as “assembled contigs” rather than MAGs. Assembly yielded a substantial number of contigs across samples, ranging from approximately 175,000 to 375,000 per sample. The distribution of contig counts was similar between the natural- and mixed-fed cohorts. Median contig counts were nearly identical between groups, and within-group variation, as shown by the interquartile range (the height of the boxes), was comparable. Statistical analysis confirmed no significant difference in contig counts between the two cohorts (*p*-value = 1.0).

### Alpha Diversity

Alpha diversity indices were used to assess microbial richness and evenness within each dietary group (Figure 3). The Observed index showed higher microbial diversity in natural feed compared to mixed feed (*p*=0.0019). The Shannon index also indicated higher diversity in natural-fed samples (*p=* 0.0274, <0.05) and showed significant differences within and between both groups. The Inverse Simpson index showed a similar trend of higher richness and evenness in the natural-fed samples, but did not reach statistical significance (*p*=0.0587). Overall, natural-fed cattle exhibited greater microbial richness and more consistent diversity than that of mixed-fed cattle.

**Figure 3.**
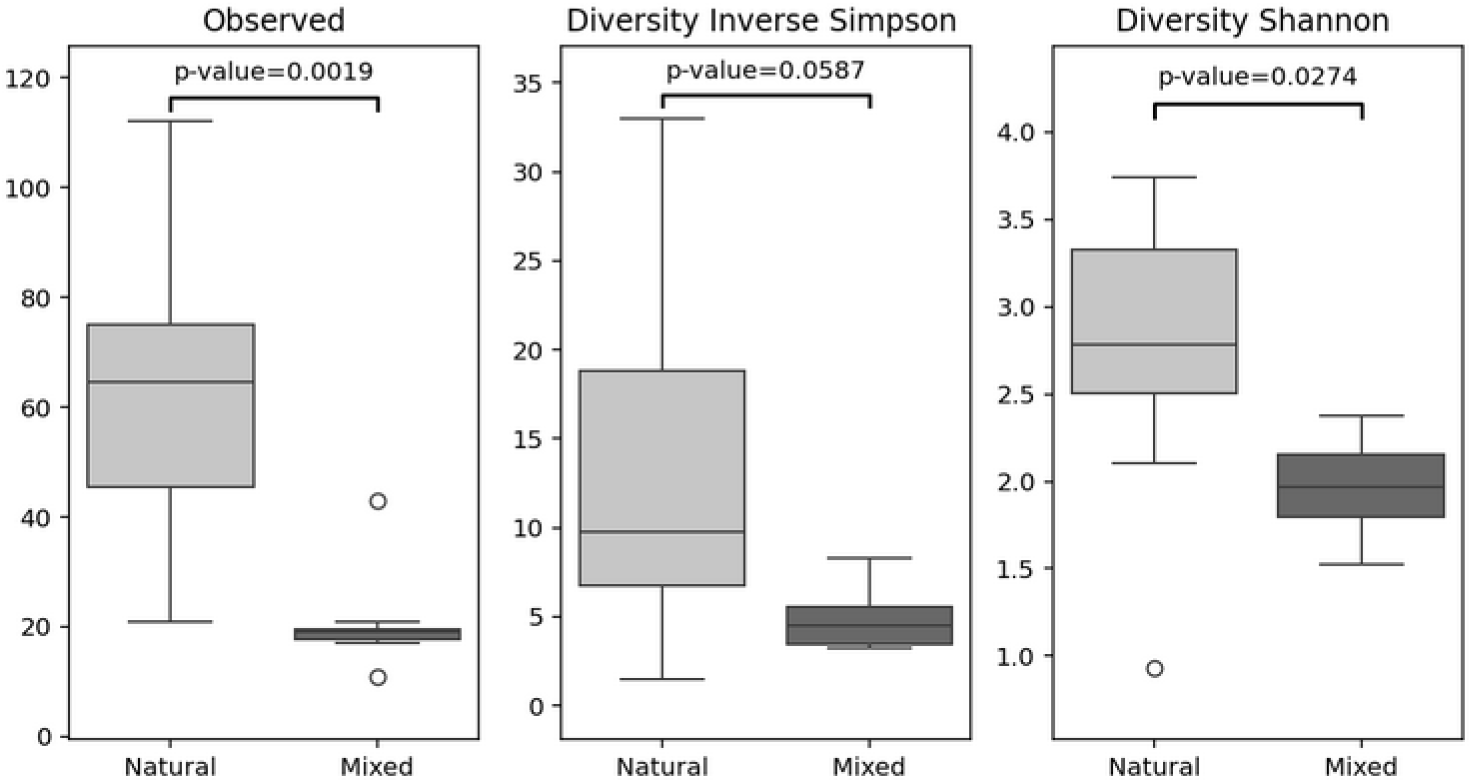
Alpha diversity of the rumen microbiome in natural- and mixed-fed cows. Boxplots show Observed richness, Inverse Simpson, and Shannon diversity indices, with *p*-values indicating significant reductions in diversity in the mixed-fed group.

### Beta Diversity

Beta diversity analysis using the Bray-Curtis dissimilarity revealed notable differences in microbial community composition between the two dietary groups (Figure 4). PERMANOVA (Permutational Multivariate Analysis of Variance) analysis showed a significant effect of diet, yielding an F-value of 1.944 and *p*-value of 0.01, explaining 19.44% of the total variation in community structure between the two groups. These results indicate that microbial community structure differs between natural- and mixed-fed cattle, and these differences are unlikely due to random variation. The separation observed in the PCoA indicates distinct microbial communities associated with each group, suggesting a diet-associated shift in community composition.

**Figure 4.**
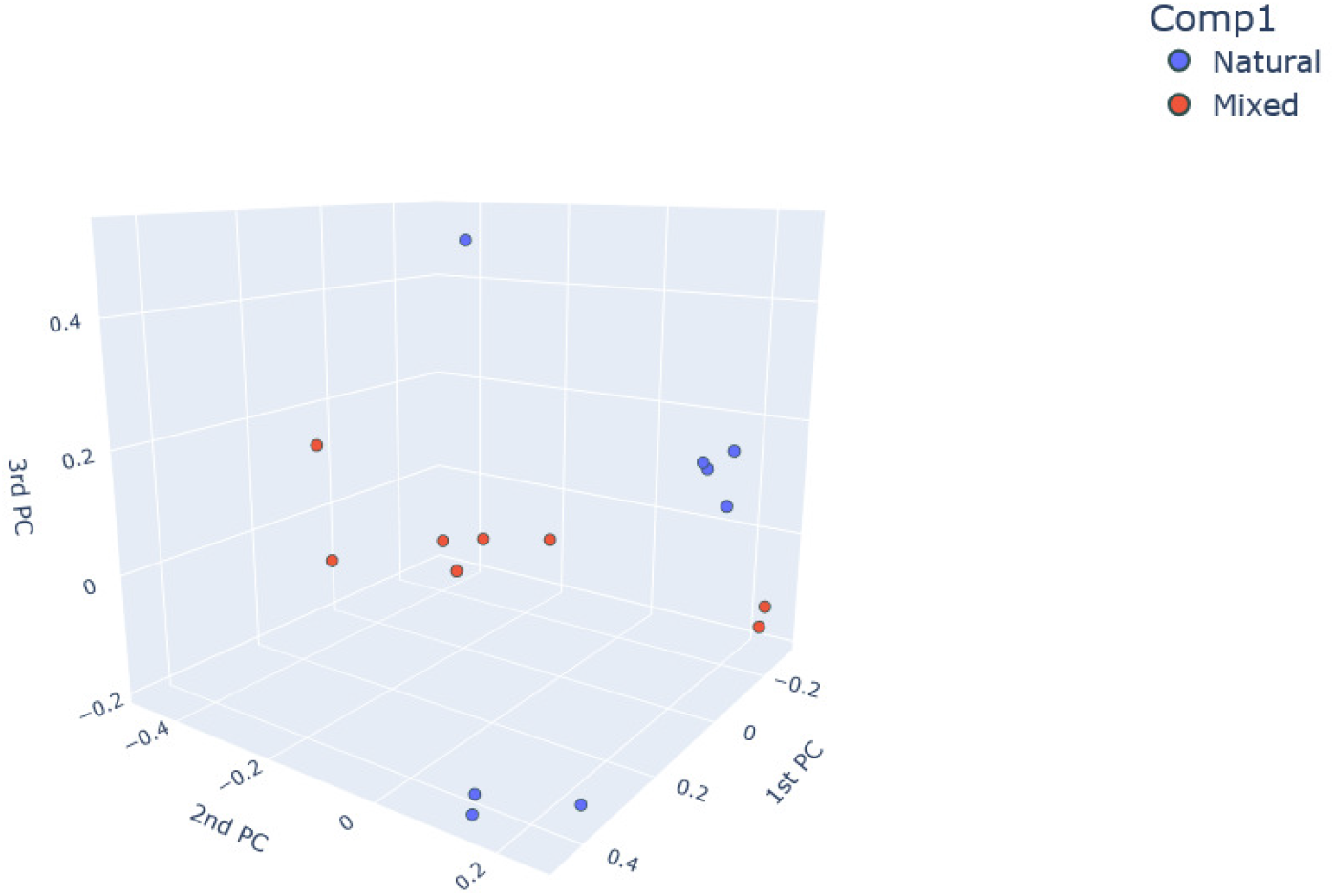
Beta diversity of rumen microbiomes from natural- and mixed-fed cows. 3D PCoA plot showing separation of samples based on community composition, illustrating diet-associated clustering patterns. The blue dots represent natural, and the red dots indicate the mixed feed samples.

### Differential Abundance Analysis

Differential abundance analysis using the ANOVA-Like Differential Expression (ALDEx2) was performed to identify taxa associated with each feeding group. ALDEx2 is a robust statistical tool designed to visualize and identify taxa as potential biomarkers on sequencing data that are differentially abundant between cows fed mixed and natural feeds. In Figure 5, the positive direction shows the effect size for the naturally fed, and the negative direction for the mixed-fed. *Oscillibacter ruminantium* showed the highest effect size and was identified as a biomarker for natural-fed group. In contrast, *Anaerotigum faecicola, Anaerovibrio slackiae, O. valericigenes*, and *Succinivibrio faecicola* were enriched in the mixed-fed samples. Among these, *Succinivibrio faecicola* and *Anaerovibrio slackiae* showed the strongest negative effect sizes and were key biomarkers of mixed-fed group.

**Figure 5.**
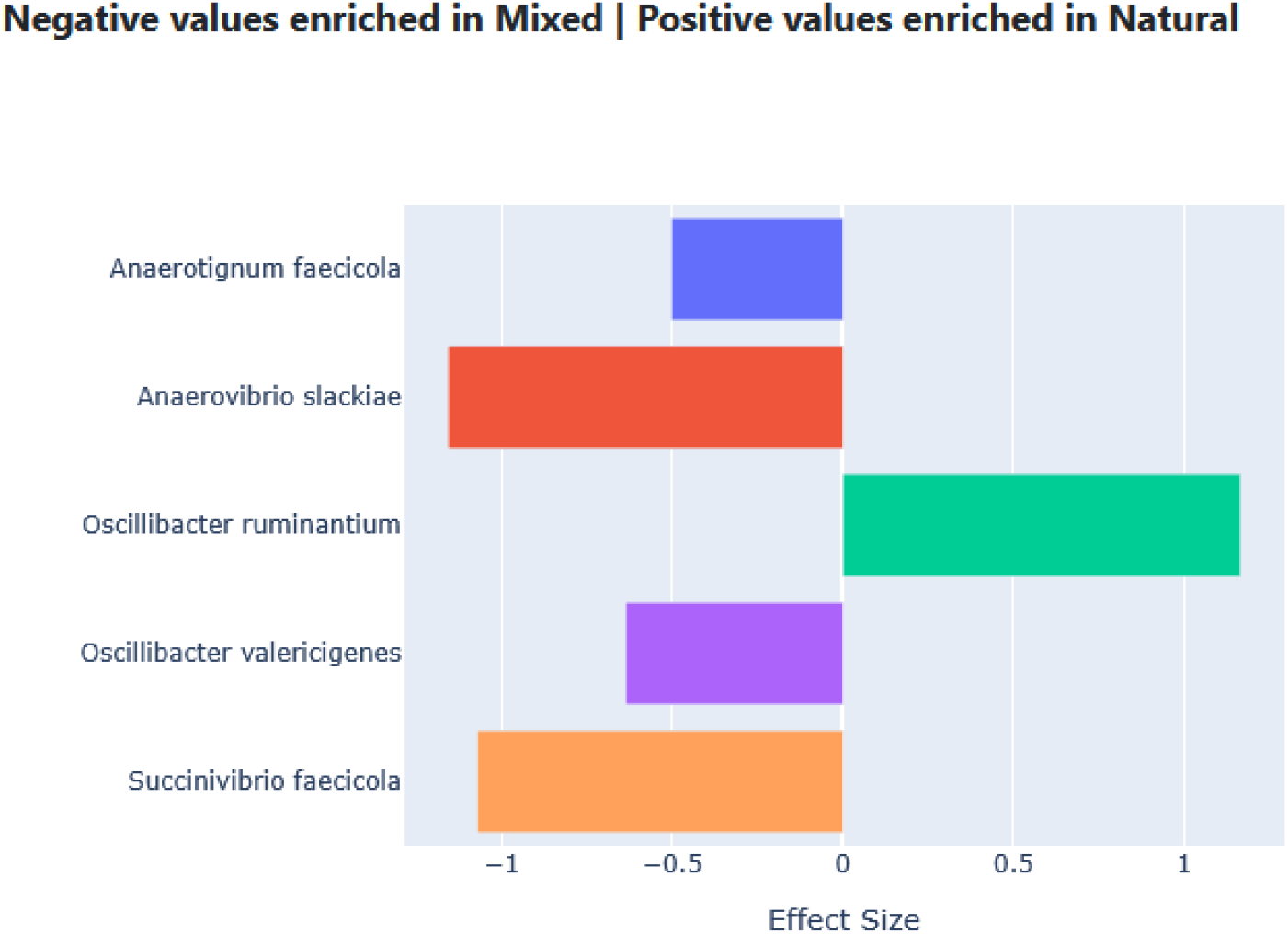
Differentially abundant taxa associated with natural and mixed feeding. LEfSe-derived effect sizes highlight microbial biomarkers, with positive values enriched in natural-fed cows and negative values enriched in mixed-fed cows.

### Antimicrobial resistance

Figure 6 presents a heatmap visualizing the count of antimicrobial resistance genes (ARGs) identified through shotgun metagenomic analysis of fecal samples from cattle subjected to different dietary regimens. The antimicrobial resistance gene analysis revealed distinct resistome profiles associated with feeding practices. A notable finding was the consistent enrichment of tetracycline resistance genes in the mixed-fed, with relatively higher abundance and uniform distributions across samples. Beta-lactam resistance genes were detected in both groups but showed variable patterns. The natural-fed group exhibited a broader range and higher peak abundance of these genes. In contrast, the mixed-fed group showed more consistent but generally lower counts. The natural-fed samples also displayed more heterogeneous resistance profiles. Vancomycin, chloramphenicol, and carbapenem resistance were detected in a single natural-fed sample (Sample-RA), while streptomycin resistance was primarily observed in natural-fed samples (Sample-3, 4, and RA) and only in one mixed-fed sample (Sample-A).

**Figure 6.**
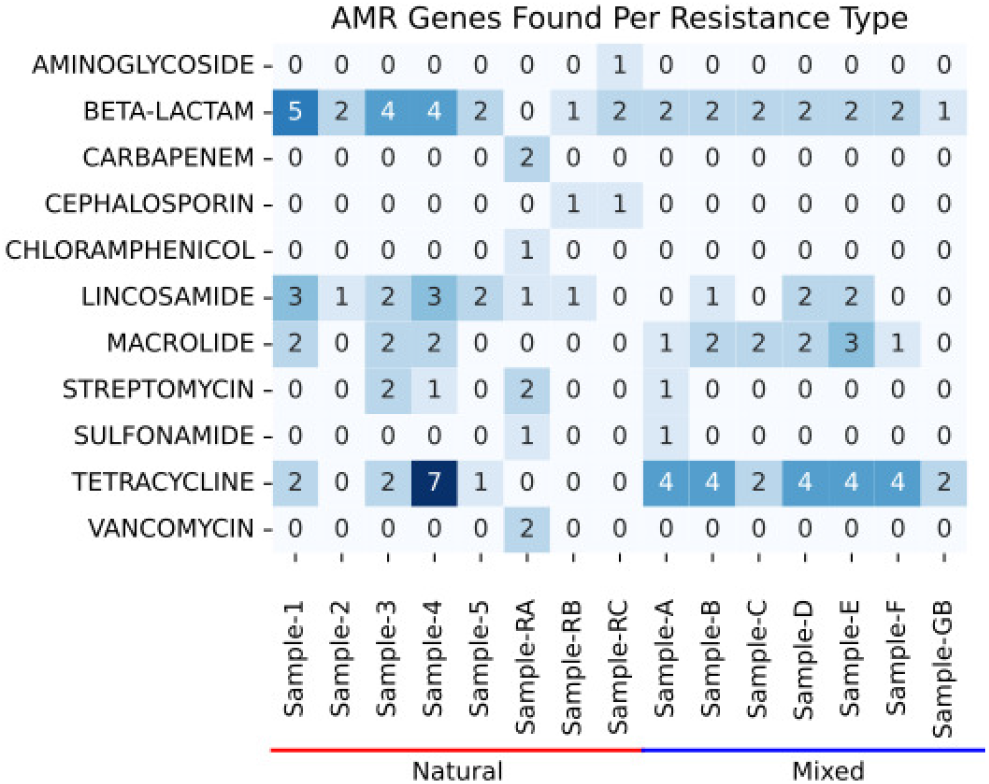
Heatmap of antimicrobial resistance gene (ARG) abundance across different feed type cohorts. The cohort for which each sample belongs is written below the red (natural) and blue (mixed) lines. The color intensity and numerical values in each cell represent the number of distinct ARGs detected for a specific resistance type within a given sample.

### Functional Diversity

Functional diversity analysis showed a consistent trend toward higher functional diversity in the natural-fed group, although these differences did not reach the conventional threshold of statistical significance (Figure 7). The number of Observed Pathways (Panel A), a measure of functional richness, was higher in the natural-fed group (*p*=0.2076). Similar trend of greater functional diversity and evenness in the natural-fed group compared to the mixed-fed group was also observed for the Inverse Simpson Index (Panel B, *p*=0.0929) and the Shannon Index (Panel C, *p*=0.0742). Notably, the natural-fed group consistently showed greater variability between samples, as evidenced by a larger interquartile range, while the mixed-fed group demonstrated a more homogeneous functional richness across samples.

**Figure 7.**
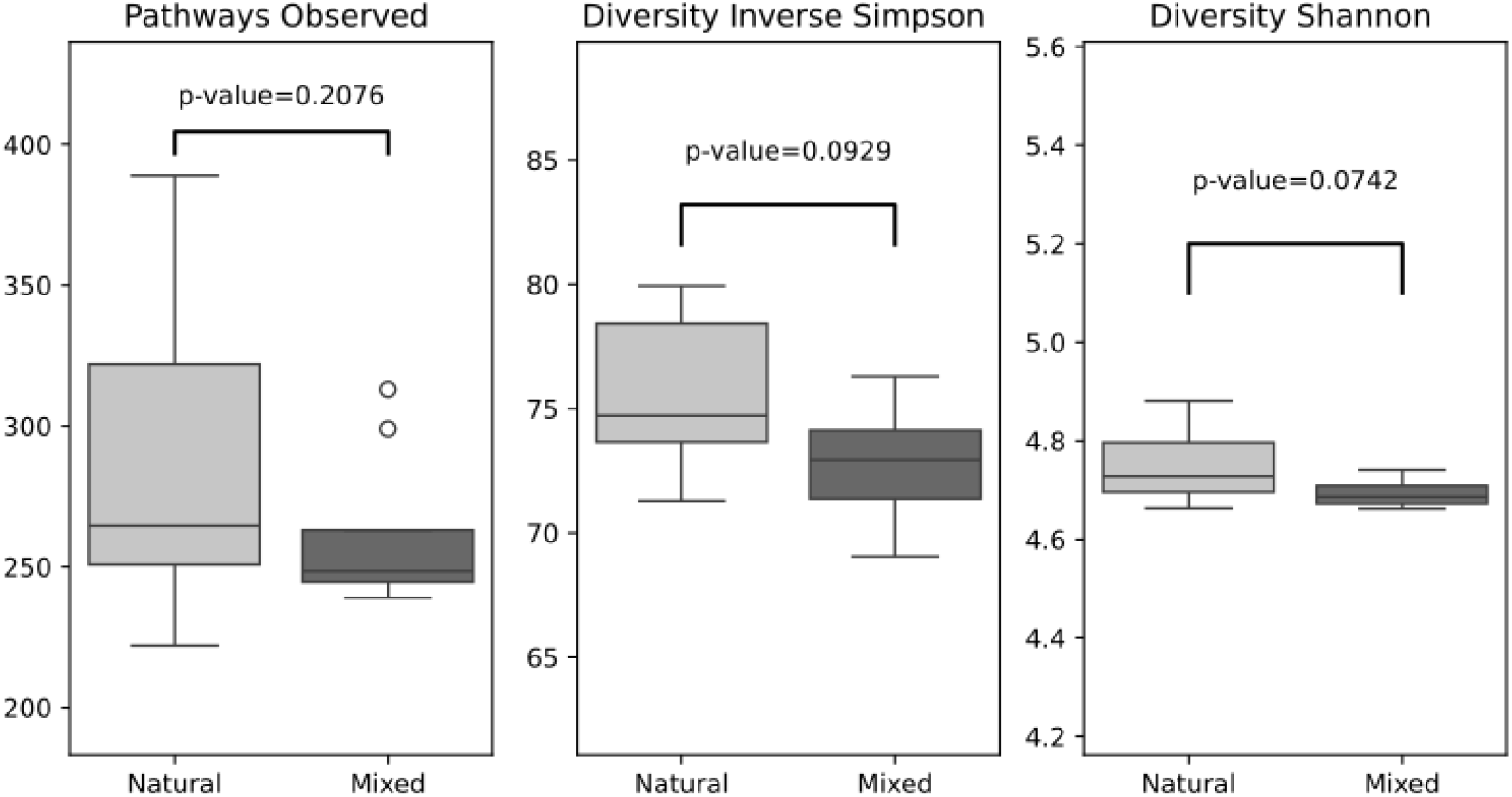
Distribution of functional diversity metrics between dietary cohorts as calculated from pathway abundance values. The three panels in the figure represent (A) the number of observed pathways, (B) the Inverse Simpson diversity index, and (C) the Shannon diversity index of functional pathways between the natural and mixed feed groups. The *p-value* from statistical comparisons for each metric is indicated.

Differential pathways analysis (Figure 8) showed distinct microbial metabolic signatures driven by diet and identified several pathways enriched in the natural-fed group, including glutathione metabolism, axon regeneration etc. Fewer pathways (i.e., bile secretion) were relatively enriched in the mixed-fed group, although these differences appear less robust.

**Figure 8.**
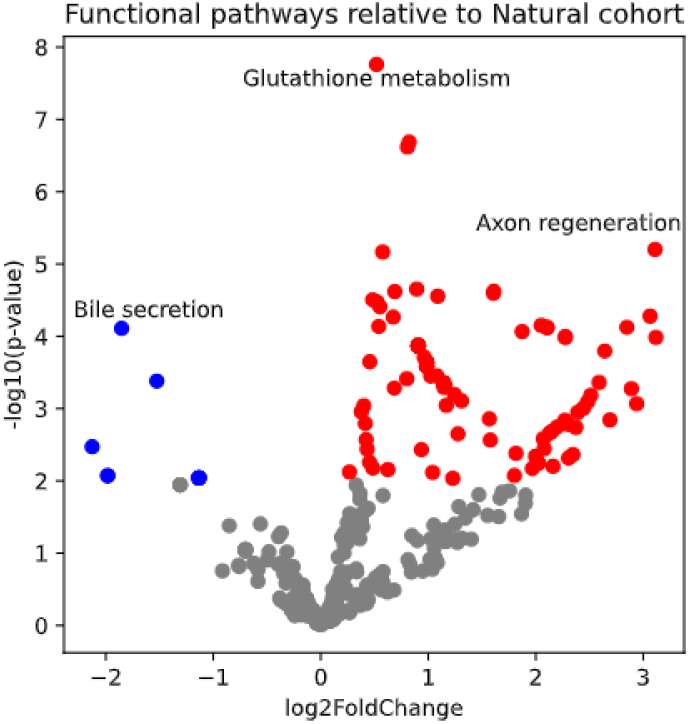
Differential abundance of microbial metabolic pathways between Natural and Mixed feed cohorts. The volcano plot displays KEGG pathways based on their log_2_ fold-change (x-axis) and statistical significance (-log_10_ *p-value*, y-axis) when comparing the Natural versus Mixed feed groups. Pathways significantly overexpressed in the Natural cohort are shown in red, while underexpressed pathways are shown in blue (*p* < 0.05). Key significantly differentiated pathways are labeled, including glutathione metabolism, which was enriched in the Natural cohort. Gray points represent pathways with no significant differential abundance between cohorts.

## Discussion

### Microbial community structure

Among the many factors that influence the rumen microbiome, diet is considered one of the strongest drivers of the microbiome community composition (Gong et al., 2020). In the present study, several bacterial phyla differed between the two feeding groups. *Actinomycetota, Bacillota, Pseudomonadota*, and *Thermodesulfobacteriota* were detected in both groups. *Bacillota* and *Thermodesulfobacteriota* were also been reported as predominant phyla in natural feeding of Kazakh bulls (Sizova et al., 2025). *Actinomycetota* has been reported to increase during ruminal acidosis, facilitating the fermentation of dietary glycans (Han et al., 2020; Modrackova et al., 2020). *Bacilliota* has been reported to be enriched in the rumen of cattle during the postweaning diet (Peraza et al., 2024). In addition, the dominance of *Bacillota* taxa improved feed conversion efficiency (Sizova et al., 2025). *Pseudomonadota* (previously known as *Proteobacteria*) has also been reported at higher abundance in the rumen of animals consuming forage-based diet (Auffret et al., 2017). The *Bacteroidota* phylum has predominantly been reported in naturally fed ruminants (Sizova et al., 2025); which is consistent with our findings in this study. *Mycoplasmatota* (known as *Tenericutes*) has been reported in the rumen of goats (Xue et al., 2022). Guo et al., 2021 reported enrichment of *Candidatus* and *Spirochaetota* in the rumen of cows with laminitis.

At the Family level, the detection of *Acholeplasmataceae* and *Acidaminococcaceae* in the natural-fed group is in agreement with previous reports from goats fed a natural diets (Belanche et al., 2023; Rebeiro et al., 2017). *Prevotellaceae* and *Oscillospiraceae* have been widely known as members of the bovine rumen microbiome. Higher abundance of *Prevotellaceae* has been reported in both Jersey cow and Holistein cow rumen samples (Paz et al., 2016). Within the family, *Prevotella* is closely associated with propionate production in the rumen (Joseph et al., 2018). Propionate is generally negatively associated with methane production (Spence et al., 2006; Bryant et al., 1958). Therefore, an increase in *Prevotella* may favor reduced methane emissions and could potentially contribute to reducing greenhouse gas emissions from ruminants. Members of the *Oscillospiraceae* are also known to contribute to short-chain fatty acid fermentation in ruminants (Akram et al., 2025).

### Microbial Diversity

Cows fed natural feed showed greater microbial variability, as reflected by the significant differences in the alpha diversity indices (Observed, Inverse Simpson, and Shannon). The rumen microbiome of natural feed-fed cows showed greater richness and evenness than that of mixed feed-fed cows. Beta diversity analysis using Bray-Curtis’s dissimilarity and PCoA analysis also revealed statistically significant clustering by feed type (PERMANOVA *p* = 0.01). This separation suggests that the two feeding regimens were associated with distinct microbial community structures. Higher diversity may reflect a more complex microbiome with greater capacity to adapt to environmental and dietary variation (Stoffel *et al*., 2020). The rumen microbiota contributes to host protection through the production of antimicrobials and by inhibiting the growth of pathogenic microorganisms (Baumler, *et al*., 2016; Lettat *et al*., 2012). A balanced rumen microbiome can also support the growth performance of the host (Shin *et al*., 2019).

Differential analysis identified *Oscillibacter ruminantium* as a strong biomarker for natural-fed groups with the highest effect size. It has previously been isolated from the rumen of Korean native cattle and has been known to digest many carbohydrates and produce butyrate as an end product (Lee *et al*. 2013). In contrast, *Anaerotigum faecicola, Anaerovibrio slackiae, O. valericigenes*, and *Succinivibrio faecicola* were enriched in the mixed-fed samples. Among these, *Succinivibrio faecicola* and *Anaerovibrio slackiae* showed strong negative effect sizes and were the biomarkers for the mixed-fed group. *Anaerovibrio sp*. are recognized rumen lipolytic bacteria, whereas *Succinivibrio faecicola* has been reported specifically from cow feces (Prins *et al*., 1975; Choi *et al*., 2022).

*Acinetobacter* has been reported to cause cellulase degradation in the rumen. When mixed with cellulase nanoparticles, *Acinetobacter pittii* has also been explored in combination with cellulase nanoparticles as part of a proof-of-concept drug delivery system in cattle (Sariboga *et al*., 2024). *Acinetobacter baumanii*, an important human pathogen, has also been detected in cattle and pig rumen-associated environments as novel clones (Estrada *et al*., 2022). *Comamonas* has been reported from the dorsal epithelium of the rumen and is considered an oxygen-scavenging bacterium that helps maintain anaerobic conditions in the epithelium and may involve in nitrogen metabolism (Li *et al*., 2024). *Aristaeella hokkaidonensis* utilizes a broad range of carbohydrates, including arabinose, galactose, glucose, xylose, cellobiose, lactose, maltose, melibiose, sucrose, and trehalose, and ferments substrates such as dextrin, pectin, starch, xylan, and salicin, with ethanol, hydrogen, acetate, and lactate as major end products (Mahoney-Kurpe *et al*., 2023). *Oscillibacter ruminantium*, isolated from Korean native cattle, utilizes D-glucose, D-ribose and D-xylose, and can use elemental sulfur, sulfate, thiosulfate, and nitrate as electron acceptors (Lee *et al*., 2013). *Flintibacter porci*, a member of the family *Oscillospiraceae*, has been reported to produce butyrate when isolated from piglet manure (Niu *et al*., 2025). Dysosmobacter is known to utilize maltose and maltodextrin, and has been linked to sugar alcohol metabolism, including enrichment of inositol catabolism within the rumen microbiome. Species of *Phascolarctobacterium* are generally asaccharolytic and use succinate as a substrate (Conteville *et al*., 2023). *Sharpea* species are known to produce lactate as a fermentation end product in the sheep rumen (Kumar *et al*., 2018).

To our knowledge, this is the first report of *Aliarcobacter* spp. in Bangladeshi cattle, following an earlier report of its presence in ready-to-eat poultry meat in Bangladesh (Mahmud *et al*., 2023). *Aliarcobacter* spp. have been described as an ‘emerging zoonotic pathogen’ (Ferreira, *et al*., 2016) and later as a ‘potential food-borne pathogen’ (Celik *et al*., 2022). Species of this genus can cause diarrhoea in both animal and humans, and farm animals may serve as reservoirs of infection (Celik *et al*., 2022). These organisms can colonize the intestines of animals with or without diarrhoea and may contaminate meat or milk through fecal shedding, thereby posing a potential public health risk.

### Functional diversity

Naturally fed cattle exhibited enrichment of several metabolic pathways, particularly glutathione metabolism and axon regeneration-related signaling, relative to mixed-fed cattle. This is generally consistent with established evidence that forage-based diets promote greater rumen microbial diversity, metabolic redundancy, and adaptation to oxidative stress (Henderson *et al*., 2015; Tapio *et al*., 2017). Glutathione plays a central role in protecting anaerobic rumen microorganisms from oxidative and nitrosative stress (Fahey and Sundquist, 1991). Natural feed sources such as grass and hay generally promote fiber fermentation, hydrogen production, and redox fluctuations, conditions under which glutathione-dependent antioxidant pathways may become critical for microbial survival (Mao *et al*., 2013; Liu *et al*., 2016). Increased glutathione-related activity has previously been linked to enhanced resilience of fibrolytic bacteria under high-fiber diets (Huws *et al*., 2018), suggesting that naturally fed cattle may harbor microbial communities better adapted to the oxidative fluctuations associated with plant-based fermentation. The higher microbial diversity observed in the present study is consistent with this interpretation

The enrichment of pathways related to axon regeneration in rumen metagenomes is consistent with the observation that many KEGG pathways originally annotated for neuronal repair overlap with microbial pathways involved in membrane remodeling, lipid metabolism, and cellular signaling (Kanehisa *et al*., 2016). Recent metagenomic studies suggest that such pathway assignments may reflect enhanced glycerophospholipid turnover, fatty acid biosynthesis, and maintenance of membrane integrity in microbial communities (Douglas *et al*., 2020). These functions are likely important for rumen microbes exposed to high-fiber diets, where adaptation to variable pH, polysaccharide load, and fermentation end-products is continuously required (Jami *et al*., 2013).

In contrast, relatively few pathways were underexpressed in the natural cohort, with bile secretion being the most notable, although statistical support was weaker. This pattern is plausible, as bile-associated pathways are more commonly enriched in microbiomes exposed to diets with higher proportions of concentrate or lipid content (Devendran *et al*., 2021). Bile acids can exert strong selective pressures on microbial communities by disrupting cell membranes and altering redox states (Wahlstrom *et al*., 2016). Mixed-fed cattle, which consumed more diverse feed types including concentrates, may have experienced greater exposure to lipid and bile-related metabolites, which could explain the relative enrichment of these pathways. However, the lower statistical support suggests that these differences were modest and may also reflect inter-animal variation.

### Antimicrobial resistance profile

The feed-associated resistome profiles identified here provide useful insights into antimicrobial resistance in livestock production systems. The consistent detection of tetracycline resistance genes across mixed-feed cows, compared to their more variable occurrence in the natural-fed cohort, suggests a more uniform resistome pattern associated with mixed feeding. Although tetracyclines may be directly used in some feed formulations, the consistency of this pattern may also reflect broader ecological effects of commercial feed, including changes in gut nutrient availability and co-selection for resistant taxa. In contrast, cattle on a natural diet exhibited a more heterogeneous resistome, including sporadic detection of resistance genes associated with high-priority drugs like carbapenems and vancomycin. This pattern may reflect more diverse environmental exposures, including soil, water, and variable forage sources, rather than a single consistent dietary input. From a practical perspective, these findings suggest an important trade-off. Commercial feeding practices may support production efficiency, but they may also favor predictable resistance risks. Conversely, natural feeding may be associated with greater microbial diversity, yet it does not eliminate exposure to environmentally derived resistance genes.

Previous studies have shown that the diversity and abundance of rumen bacteria are closely linked to feed composition and type (Carberry *et al*., 2012). Our study similarly revealed clear differences between the microbiomes of cows fed natural and mixed feeds. It has also been reported that highly nutritious feeds can reduce rumen microbial diversity and may contribute to subacute ruminal acidosis (SARA), which is detrimental to cattle health (Bevans *et al*., 2005). However, both cohorts in the present study appeared to maintain relatively stable rumen ecosystems. Cows fed natural diets showed higher microbial richness, which may favor beneficial rumen function. In contrast, the mixed-fed group maintained a stable but functionally distinct microbiome, likely adapted to a higher energy diet. These findings are consistent with the idea that the ruminal microbial composition can be modulated through feed formulation to improve microbial stability and support animal production (Russell and Rychlik, 2001). Adjusting the ratio of concentrate to roughage in feeds may help optimize productivity, reduce energy loss, and decrease methane emissions (Makkar and Beever, 2013). In Bangladesh, such feed management strategies may help support cattle health, improve production efficiency, and reduce environmental impact, while also contributing to better antimicrobial stewardship. Overall, the metabolic pathways observed in this study reinforce the concept that diet composition is a primary driver of rumen microbial function. Natural forage-based diets promoted oxidative stress adaptation, membrane remodeling, and metabolic flexibility, while mixed diets shifted functional emphasis toward pathways consistent with higher lipid and concentrate intake.

Several limitations should be acknowledged. As a pilot study, the modest sample size (n=16) and regional scope may not capture the full variability within each feed type and be limit generalization to other production settings and/or regions. A multi-center study with a larger sample size is recommended for statistical robustness. The study also did not account for temporal variations in microbiome composition, highlighting the need for longitudinal sampling. In addition, resistome profiling from metagenomic data reflects genetic potential rather than confirmed gene expression or phenotype. Future transcriptomic and phenotypic analyses for validation. Despite these limitations, this study shows that feeding practice is closely associated with rumen microbiome composition, diversity, metabolic function, and antimicrobial resistance patterns in Bangladeshi cattle. Natural grass-based diets were associated with higher microbial richness, distinct taxa, and pathways linked to adaptive microbial functions, emphasizing the importance of feeding strategies in rumen health, productivity, and antibiotic stewardship. Longer-term studies will be important to determine how sustained dietary patterns influence rumen microbial stability, potential dysbiosis, and downstream effects on cattle productivity and health in Bangladesh.

## Data availability

The data has been submitted in the National Centre for Biotechnology Institute. The BioProject is identified by PRJNA1355225.

## Funding declaration

The research was funded by the Centennial Research Grant conferred by the University of Dhaka, Bangladesh.

## Competing interest

The authors declare that there are no competing interests.

## Author contribution

S.A. conceptualized the research, managed the fund and wrote the manuscript. N.A.H. wrote the manuscript and did the analysis. M.J.N. wrote the manuscript and did the analysis. M.C. contributed to writing the manuscript. F.N. wrote part of the manuscript and helped with laboratory work. E.M. helped with labwork.

## Ethics declaration

Not applicable.

## References

1. Eisler, M.C., Lee, M.R., Tarlton, J.F., Martin, G.B., Beddington, J., Dungait, J.A., et al. (2014). Agriculture: steps to sustainable livestock. Nature. 507, 32–34.

2. O’Hara, E., Neves, A.L.A., Song, Y., and Guan, L.L. (2020). The role of the gut microbiome in cattle production and health: driver or passenger? Annu. Rev. Anim. Biosci. 8, 199–220.

3. Mao, S., Zhang, M., Liu, J., and Zhu, W. (2015). Characterizing the bacterial microbiota across the gastrointestinal tracts of dairy cattle: membership and potential function. Sci. Rep. 5, 16116.

4. Stewart, R.D., Auffret, M.D., Warr, A., Walker, A.W., Roehe, R., and Watson, M. (2019). Compendium of 4,941 rumen metagenome-assembled genomes for rumen microbiome biology and enzyme discovery. Nat. Biotechnol. 37, 953–961.

5. Alberdi, A., Andersen, S.B., Limborg, M.T., Dunn, R.R., and Gilbert, M.T.P. (2021). Disentangling host-microbiota complexity through hologenomics. Nat. Rev. Genet. 22, 148–159.

6. Eastridge, M. (2006). Major advances in applied dairy cattle nutrition. J. Dairy Sci. 89, 1311–1323.

7. Zhao, C., Hu, X., Zhang, N., and Fu, Y. (2025). Emerging role of ruminal microbiota in the development of perinatal bovine diseases. Anim. Zoonoses 1, 86–98.

8. Petra, L., Hold, G.L., and Flint, H.J. (2014). The gut microbiota, bacterial metabolites and colorectal cancer. Nat. Rev. Microbiol. 12, 661–672.

9. Martínez-Álvaro, M., Auffret, M.D., Stewart, R.D., Dewhurst, R.J., Duthie, C-A., Rooke, J.A., et al. (2020). Identification of complex rumen microbiome interaction within diverse functional niches as mechanisms affecting the variation of methane emissions in bovine. Front. Microbiol. 11, 659.

10. Artzi, L., Bayer, E.A., and Morais, S. (2017). Cellulosomes: bacterial nanomachines for dismantling plant polysaccharides. Nat. Rev. Microbiol. 15, 83–95.

11. Seshadri, R., Leahy, S.C., Attwood, G.T., Teh, K.H., Lambie, S.C., Cookson, A.L., et al. (2018). Cultivation and sequencing of rumen microbiome members from the Hungate1000 Collection. Nat. Biotechnol. 36, 359–367.

12. Chae, T.U., Ahn, J.H., Ko, Y-S., Kim, J.W., Lee, J.A., Lee, E.H., et al. (2020). Metabolic engineering for the production of dicarboxylic acids and diamines. Metab. Eng. 58, 2–16.

13. Dimroth, P., and Schink, B. (1998). Energy conservation in the decarboxylation of dicarboxylic acids by fermenting bacteria. Arch. Microbiol. 170, 69–77.

14. Leahy, S.C., Janssen, P.H., Attwood, G.T., Mackie, R.I., McAllister, T.A., and Kelly, J.W. (2022). Electron flow: key to mitigating ruminant methanogenesis. Trends Microbiol. 30, 209–212.

15. Cholewinska, P., Czyz, K., Nowakowski, P., and Wyrostek, A. (2020). The microbiome of the digestive system of ruminants - a review. Anim. Health Res. Rev. 21, 3–14.

16. Ransom-Jones, E., Jones, D.L., McCarthy, A.J., and McDonald, J.E. (2012). The Fibrobacteres: an important phylum of cellulose-degrading bacteria. Microb. Ecol. 63, 267–281.

17. Driks, A. (2003). The dynamic spore. Proc. Natl. Acad. Sci. U. S. A. 100, 3007–3009.

18. Liggenstoffer, A.S., Youssef, N.H., Couger, M.B., and Elshahed, M.S. (2010). Phylogenetic diversity and community structure of anaerobic gut fungi (phylum Neocallimastigomycota) in ruminant and nonruminant herbivores. ISME J. 4, 1225.

19. Tropini, C., Earle, K.A., Huang, K.C., and Sonnenburg, J.L. (2017). The gut microbiome: connecting spatial organization to function. Cell Host Microbe 21, 433–442.

20. Borrel, G., Adam, P.S., and Gribaldo, S. (2016). Methanogenesis and the Wood-Ljungdahl pathway: an ancient, versatile, and fragile association. Genome Biol. Evol. 8, 1706–1711.

21. Donaldson, G.P., Lee, S.M., and Mazmanian, S.K. (2016). Gut biogeography of the bacterial microbiota. Nat. Rev. Microbiol. 14, 20–32.

22. Kamke, J., Kittelmann, S., Soni, P., Li, Y., Tavendale, M., Ganesh, S., et al. (2016). Rumen metagenome and metatranscriptome analyses of low methane yield sheep reveals a Sharpea-enriched microbiome characterized by lactic acid formation and utilization. Microbiome. 4, 56.

23. Huque, K., and Sarker, N. (2014). Feeds and feeding of livestock in Bangladesh: performance, constraints and options forward. Bangladesh J. Animal Sci. 43, 1–10.

24. Kirbis, A. (2007). Microbiological screening method for detection of aminoglycosides, β-lactams, macrolides, tetracyclines and quinolones in meat samples. Slov. Vet. Res. 44, 11–18.

25. Lin, L., Xie, F., Sun, D., Liu, J., Zhu, W., and Mao, S. (2019). Ruminal microbiome-host crosstalk stimulates the development of the ruminal epithelium in a lamb model. Microbiome 7, 83.

26. Li, H., Handsaker, B., Wysoker, A., Fennell, T., Ruan, J., Homer, N., and Durbin, R. (2009). The Sequence Alignment/Map format and SAMtools. Bioinformatics 25, 2078–2079.

27. Buchfink, B., Xie, C., and Huson, D.H. (2014). Fast and sensitive protein alignment using DIAMOND. Nature Methods 12, 59–60.

28. Kanehisa, M., Furumichi, M., Tanabe, M., Sato, Y., and Morishima, K. (2017). KEGG: new perspectives on genomes, pathways, diseases and drugs. Nucleic Acids Res. 45, D353–D361.

29. Ye, Y., and Doak, T.G. (2009). A parsimony approach to biological pathway reconstruction/inference for genomes and metagenomes. PLoS Comput. Biol. 5, 1–8.

30. Langmead, B., and Salzberg, S.L. (2012). Fast gapped-read alignment with Bowtie 2. Nature Methods. 9, 357–359.

31. Yoon, S., Ha, S., Kwon, S., Lim, J., Kim, Y., Seo, H., and Chun, J. (2019). Introducing EzBioCloud: a taxonomically united database of 16S rRNA gene sequences and whole-genome assemblies. Int. J. Syst. Evol. Microbiol. 67, 1613–1617.

32. Shannon, C.E. (1948). A mathematical theory of communication. Bell Syst. Tech. J. 27, 379–423.

33. Simpson, E.H. (1949). Measurement of diversity. Nature. 163, 688.

34. Bray, J.R., and Curtis, J.T. (1957). An ordination of the upland forest communities of Southern Wisconsin. Ecol. Monogr. 27, 325–349.

35. Benjamini, Y., and Hochberg, Y. (1995). Controlling the false discovery rate: a practical and powerful approach to multiple testing. J. R. Stat. Soc. Ser. B Methodol. 57, 289–300.

36. Love, M.I., Huber, W., and Anders, S. (2014). Moderated estimation of fold change and dispersion for RNA-seq data with DESeq2. Genome Biol. 15, 550.

37. Gong, G., Zhou, S., Luo, R., Gesang, Z., and Suolang, S. (2020). Metagenomic insights into the diversity of carbohydrate-degrading enzymes in the yak fecal microbial community. BMC Microbiol. 20, 302.

38. Sizova, E., Yausheva, E., Miroshnikov, S., Kamirova, A., and Shoshin, D. (2025). Ruminal digestion, gastrointestinal microbial profile, and metabolic pathways after the introduction of silicon-containing ultrafine particles into bull. Vet. World 18, 1070–1081.

39. Han, Z., Li, A., Pei, L., Li, K., Jin, T., Li, F., et al. (2020). Milk replacer supplementation ameliorates growth performance and rumen microbiota of early-weaning Yimeng black goats. Front. Vet. Sci. 7, 572064.

40. Modrackova, N., Vlkova, E., Tejnecky, V., Schwab, C., and Neuzil-Bunesova, V. (2020). Bifidobacterium β-glucosidase activity and fermentation of dietary plant glucosides is species and strain specific. Microorganisms. 8, 839.

41. Peraza, P., Fernández-Calero, T., Naya, H., Sotelo-Silveira, J., and Navajas, E.A. (2024). Exploring the linkage between ruminal microbial communities on postweaning and finishing diets and their relation to residual feed intake in beef cattle. Microorganisms. 12, 2437.

42. Auffret, M.D., Dewhurst, R.J., Duthie, C.A., et al. (2017). The rumen microbiome as a reservoir of antimicrobial resistance and pathogenicity genes is directly affected by diet in beef cattle. Microbiome. 5, 159.

43. Xue, B., Wu, M., Yue, S., Hu, A., Li, X., Hong, Q., et al. (2022). Changes in rumen bacterial community induced by the dietary physically effective neutral detergent fiber levels in goat diets. Front. Microbiol. 13, 820509.

44. Guo, J., Mu, R., Li, S., Zhang, N., Fu, Y., and Hu, X. (2021). Characterization of the bacterial community of rumen in dairy cows with laminitis. Genes. 12, 1996.

45. Belanche, A., Palma-Hidalgo, J.M., Jiménez, E., and Yáñez-Ruiz, D.R. (2023). Enhancing rumen microbial diversity and its impact on energy and protein metabolism in forage-fed goats. Front. Vet. Sci. 10, 1272835.

46. Ribeiro, G.O., Oss, D.B., He, Z., et al. (2017). Repeated inoculation of cattle rumen with bison rumen contents alters the rumen microbiome and improves nitrogen digestibility in cattle. Sci. Rep. 7, 1276.

47. Paz, H.A., Anderson, C.L., Muller, M.J., Kononoff, P.J., and Fernando, S.C. (2016). Rumen bacterial community composition in Holstein and Jersey cows is different under same dietary condition and is not affected by sampling method. Front. Microbiol. 7, 1206.

48. Joseph, R.C., Kim, N.M., and Sandoval, N.R. (2018). Recent developments of the synthetic biology toolkit for Clostridium. Front. Microbiol. 9, 154.

49. Spence, C., Wells, W.G., and Smith, C.J. (2006). Characterization of the primary starch utilization operon in the obligate anaerobe Bacteroides fragilis: regulation by carbon source and oxygen. J. Bacteriol. 188, 4663–4672.

50. Bryant, M.P., Small, N., Bouma, C., and Chu, H. (1958). Bacteroides ruminicola n. sp. and Succinimonas amylolytica; the new genus and species; species of succinic acid-producing anaerobic bacteria of the bovine rumen. J. Bacteriol. 76, 15–23.

51. Akram, A., Shahin, F., Asif, I., Bilal, A., Abbas, K., and Younas, E. (2025). Exploring the role of gut bacteria in digestive system of cow. J. Med. Health Sci. Rev. 2, 89–94.

52. Stoffel, M.A., et al. (2020). Early sexual dimorphism in the developing gut microbiome of northern elephant seals. Mol. Ecol. 29, 2109–2122.

53. Baumler, A.J., et al. (2016). Interactions between the microbiota and pathogenic bacteria in the gut. Nature. 535, 85–93.

54. Lettat, A., Nozière, P., Silberberg, M., Morgavi, D.P., Berger, C., and Martin, C. (2012). Rumen microbial and fermentation characteristics are affected differently by bacterial probiotic supplementation during induced lactic and subacute acidosis in sheep. BMC Microbiol. 12, 1–12.

55. Shin, D., Chang, S.Y., Bogere, P., Won, K.H., Choi, J.Y., Choi, Y.J., et al. (2019). Beneficial roles of probiotics on the modulation of gut microbiota and immune response in pigs. PLoS ONE. 14, e0220843.

56. Lee, G.H., Rhee, M.S., Chang, D.H., Lee, J., Kim, S., Yoon, M.H., and Kim, B.C. (2013). Oscillibacter ruminantium sp. nov., isolated from the rumen of Korean native cattle. Int. J. Syst. Evol. Microbiol. 63, 1942–1946.

57. Prins, R.A., Lankhorst, A., van der Meer, P., and Van Nevel, C.J. (1975). Some characteristics of Anaerovibrio lipolytica a rumen lipolytic organism. Antonie Van Leeuwenhoek. 41, 1–11.

58. Choi, J.Y., Park, J.E., Choi, S.H., Kim, J.S., Lee, J.S., Lee, J.H., et al. (2022). Succinivibrio faecicola sp. nov., isolated from cow faeces. Int. J. Syst. Evol. Microbiol. 72, 005631.

59. Sariboga, R., and Sarioglu, O.F. (2024). Cellulolytic characterization of the rumen-isolated Acinetobacter pittii ROBY and design of a potential controlled-release drug delivery system. Eng. Microbiol. 4, 100164.

60. Mateo-Estrada, V., Vali, L., Hamouda, A., Evans, B.A., and Castillo-Ramírez, S. (2022). Acinetobacter baumannii sampled from cattle and pigs represent novel clones. Microbiol. Spectr. 10, e0128922.

61. Çelik, C., Pınar, O., and Sipahi, N. (2022). The prevalence of Aliarcobacter species in the fecal microbiota of farm animals and potential effective agents for their treatment: a review of the past decade. Microorganisms. 10, 2430.

62. Li, S., Mu, R., Zhu, Y., Zhao, F., Qiu, Q., Si, H., et al. (2024). Shifts in the microbial community and metabolome in rumen ecological niches during antler growth. Comput. Struct. Biotechnol. J. 23, 1608–1618.

63. Mahoney-Kurpe, S.C., Palevich, N., Noel, S.J., Gagic, D., Biggs, P.J., Soni, P., et al. (2023). Aristaeella hokkaidonensis gen. nov. sp. nov. and 1Aristaeella lactis sp. nov., two rumen bacterial species of a novel proposed family, Aristaeellaceae fam. nov. Int. J. Syst. Evol. Microbiol. 73, 005831.

64. Niu, H.Y., Zhang, J., Huang, H.J., Sun, X.W., Chen, H.Y., Wang, X.M., et al. (2025). Flavonifractor porci sp. nov. and Flintibacter porci sp. nov., two novel butyrate-producing bacteria of the family Oscillospiraceae. Int. J. Syst. Evol. Microbiol. 75, 006767.

65. Conteville, L.C., da Silva, J.V., Andrade, B.G.N., Cardoso, T.F., Bruscadin, J.J., de Oliveira, P.S.N., et al. (2023). Rumen and fecal microbiomes are related to diet and production traits in Bos indicus beef cattle. Front. Microbiol. 14, 1282851.

66. Kumar, S., Treloar, B.P., Teh, K.H., McKenzie, C.M., Henderson, G., Attwood, G.T., et al. (2018). Sharpea and Kandleria are lactic acid producing rumen bacteria that do not change their fermentation products when co-cultured with a methanogen. Anaerobe. 54, 31–38.

67. Mahmud, M.M., Kabir, A., Hossain, M.Z., Mim, S.J., Yeva, I.J., Khatun, M., et al. (2023). First report of Aliarcobacter cryaerophilus in ready-to-cook chicken meat samples from super shops in Bangladesh. J. Adv. Vet. Anim. Res. 10, 113–117.

68. Ferreira, S., Queiroz, J.A., Oleastro, M., and Domingues, F.C. (2016). Insights in the pathogenesis and resistance of Arcobacter: a review. Crit. Rev. Microbiol. 42, 364–383.

69. Henderson, G., Cox, F., Ganesh, S., Jonker, A., Young, W., and Janssen, P.H. (2015). Rumen microbial community composition varies with diet and host. Nat. Microbiol. 1, 16002.

70. Tapio, I., Snelling, T.J., Strozzi, F., and Wallace, R.J. (2017). The ruminal microbiome associated with methane emissions from ruminant livestock. J. Anim. Sci. Biotechnol. 8, 7.

71. Fahey, R.C., and Sundquist, A.R. (1991). Evolution of glutathione metabolism. Adv. Enzymol. Relat. Areas Mol. Biol. 64, 1–53.

72. Mao, S.Y., Zhang, R.Y., Wang, D.S., and Zhu, W.Y. (2013). Impact of subacute ruminal acidosis (SARA) adaptation on rumen microbiota in dairy cattle. BMC Vet. Res. 9, 29.

73. Liu, J., Zhang, M., Xue, C., Zhu, W., and Mao, S. (2016). Characterization of microbial communities and predicted metabolic functions in the rumen of dairy cows fed forage- or concentrate-rich diets. Front. Microbiol. 8, 749.

74. Huws, S.A., Creevey, C.J., Oyama, L.B., et al. (2018). Addressing global ruminant agricultural challenges through understanding the rumen microbiome. Front. Microbiol. 9, 2161.

75. Kanehisa, M., Sato, Y., Kawashima, M., Furumichi, M., and Tanabe, M. (2016). KEGG as a reference resource for gene and protein annotation. Nucleic Acids Res. 44, D457–D462.

76. Douglas, G.M., Maffei, V.J., Zaneveld, J., et al. (2020). PICRUSt2 for prediction of metagenome functions. Nat. Biotechnol. 38, 685–688.

77. Jami, E., Israel, A., Kotser, A., and Mizrahi, I. (2013). Exploring the bovine rumen bacterial community from birth to adulthood. ISME J. 7, 1069–1079.

78. Devendran, S., Shrestha, R., Alves, J.M.P., Wolf, P.G., Ly, L., Hernandez, A.G., et al. (2019). Clostridium scindens ATCC 35704: integration of nutritional requirements, the complete genome sequence, and global transcriptional responses to bile acids. Appl. Environ. Microbiol. 85, e00052–19.

79. Wahlstrom, A., Sayin, S.I., Marschall, H.U., and Bäckhed, F. (2016). Intestinal crosstalk between bile acids and microbiota and its impact on host metabolism. Cell Metab. 24, 41–50.

80. Carberry, C.A., Kenny, D.A., Han, S., McCabe, M.S., and Waters, S.M. (2012). Effect of phenotypic residual feed intake and dietary forage content on the rumen microbial community of beef cattle. Appl. Environ. Microbiol. 78, 4949–4958.

81. Bevans, D.W., Beauchemin, K.A., Schwartzkopf-Genswein, K.S., McKinnon, J.J., and McAllister, T.A. (2005). Effect of rapid or gradual grain adaptation on subacute acidosis and feed intake by feedlot cattle. J. Anim. Sci. 83, 1116–1132.

82. Russell, J.B., and Rychlik, J.L. (2001). Factors that alter rumen microbial ecology. Science. 292, 1119–1122.

83. Makkar, H.P.S., and Beever, D. (2013). Optimization of feed use efficiency in ruminant production systems. FAO Animal Production and Health Proceedings 16, 1–12.

